# Candidate imprinting control regions in dog genome

**DOI:** 10.1101/2024.12.23.629822

**Authors:** Phillip Wyss, Minou Bina

**Affiliations:** Department of Chemistry, Purdue University, West Lafayette, IN 47907 USA

## Abstract

In mammals, genomic imprinting restricts the expression of a subset of genes from one of the two parental alleles. The process is regulated by imprinting control regions (ICRs) and gDMRs (germline Differentially Methylated Regions) dispersed across autosomal chromosomal DNA. An unresolved question is how to discover candidate ICRs in canine DNA genome-wide. Previously, bioinformatics analyses found a significant fraction of well-known ICRs/gDMRs in mouse, human, and *Bos taurus*. Analyses were based on finding the genomic positions of clusters of several CpG-rich motifs known as ZFBS-morph overlaps. These motifs are composite DNA elements. For this report, we performed similar studies to pinpoint candidate ICRs in the dog genome. A key feature of the bioinformatics strategy is creating density plots to mark cluster positions. In genome-wide analyses, peaks in plots effectively discovered candidate ICRs along chromosomal DNA sequences of *Canis lupus familiaris* breed Boxer. With respect to Non-Dog RefSeq Genes, several candidate ICRs are in regions analogous to ICR positions in mouse DNA, in human DNA, or both. In Boxer genome, examples include candidate ICRs for parent-of-origin-specific expression of the *MEST* isoform *PEG1, INPP5F_V2*, the *PLAGL1* isoform *ZAC1, IGF2R, PEG3*, and *GNAS* loci. In mouse, imprinted genes in these loci play important roles in developmental and physiological processes.

## INTRODUCTION

In genomic era, Man’s Best Friend offers the opportunity to identify genes that impact development, morphology, and behavior [1]. Studies of genomic imprinting would contribute to annotation and uncovering regulatory landscapes of canids. Genomic imprinting is an evolutionary and adaptive process [2]. It manifests broadly: at the level of specific cells, an organ, or a whole organism [2]. With existing drafts of Boxer genome [3], it is possible to study genomic imprinting in *Canis familiaris*. In mammals, genomic imprinting has evolved for regulating gene dosage [4, 5]. A reduction in gene dosage results when DNA methyltransferases modify the CpGs in ICRs in the maternally or paternally inherited allele [6, 7]. ZFP57 binds CpG modified hexameric sites (TGC ^me^CGC) to protect the DNA from demethylation [8-10]. Additional steps include recruitment of SETDB1 to trimethylate lysine 9 in histone H3, and HP1-mediated heterochromatin formation to completely silence imprinted genes [6].

A bioinformatics strategy demonstrated that unmodified hexameric (TGCCGC) sites occurred frequently along mouse chromosome 7 and deduced that in ICRs, additional nucleotides were required for initiating and spreading CpG methylation [11]. This hypothesis led to the discovery of a set of CpG-rich composite DNA elements known as ZFBS-morph overlaps [11, 12]. These DNA elements consist of the nonmethylated hexameric site (TGCCGC) overlapping a subset of sequences that bind the MLL1 binding units known as MLL1 morphemes [12-14]. In nucleosomes, members of the MLL1 family modify lysine 4 on histone H3 [15]. H3K4me3 marks produce chromatin states that are either transcriptionally active or poised for expression [16]. Despite the importance of genomic imprinting in placental mammals, not much is known about imprinted genes in canines, and even less about gDMRs/ICRs and their impact on embryogenesis in domesticated dogs and their wild ancestors.

## RESULTS

### Overview

In studies of Boxer genome, bioinformatics analyses produced two datasets. By uploading the datasets onto the UCSC genome browser, users create custom tracks to view the positions of TGCCGC hexamer, ZFBS-morph overlaps, and peaks in density plots. Peaks mark the positions of candidate ICRs. Typically, robust peaks cover three or more of the composite DNA elements [17]; peaks that cover two could be true or false positives. Since the genome browser is a general bioinformatics resource, it facilitates inspecting user-created tracks with respect to genomic landmarks, including annotated genes, CpG islands, and Non-Dog RefSeq Genes displayed in full, pack, or dense formats [18]. Even though the dog genome has a remarkably large number of CpG islands [19], in density plots, robust peaks are often resolved. From this outcome, one could infer that candidate ICRs correspond to rare events in dog genomic DNA. In nearly all cases, peaks appeared in unannotated Boxer DNA. Therefore, peak positions were checked with respect to Non-Dog RefSeq Genes.

### A candidate ICR in deduced *MEST* locus in Boxer

During mouse development, *Mest* is expressed biallelically in mesodermal derivatives and the hypothalamus [20, 21]. In mouse, the *Mest* locus encompasses an intragenic promoter for regulating parent-of-origin-specific expression of Peg*1*, the *Mest* imprinted isoform, and a promoter for driving transcription of biallelically-expressed gene [21]. In addition to the isoform-specific *MEST* transcript, human locus encompasses another imprinted gene (*MESTIT1*) that specifies a noncoding antisense RNA [22, 23].

With respect to human RefSeq genes, Boxer genome includes three transcripts annotated as Homo MEST, and a transcript marked as Homo MESTIT1 (Fig. 1). In closeup views, two of the *MEST* transcripts are within the intron of a longer *MEST* transcript. Keeping in mind reported studies [22-24], for Boxer, one could infer the following scenarios. The longest annotated *MEST* transcript is biallelically expressed, the shorter ones are imprinted transcripts (Fig. 1). The bioinformatics strategy predicts a candidate ICR that agrees with both scenarios. Specifically, in Boxer DNA, the density plots include a peak (a candidate ICR) in the intron of the annotated long *MEST* transcript. This intronic peak is in a CpG island (CpG174). It encompasses the transcription start sites (TSSs) of the shorter *MEST* transcripts (Fig. 1). Near the peak is the TSS of annotated MESTIT*1*. Therefore, the strategy predicted a candidate ICR for parent-of-origin-specific expression of both *PEG1* and MESTIT*1* in canine DNA (Fig. 1).

**Figure 1.**
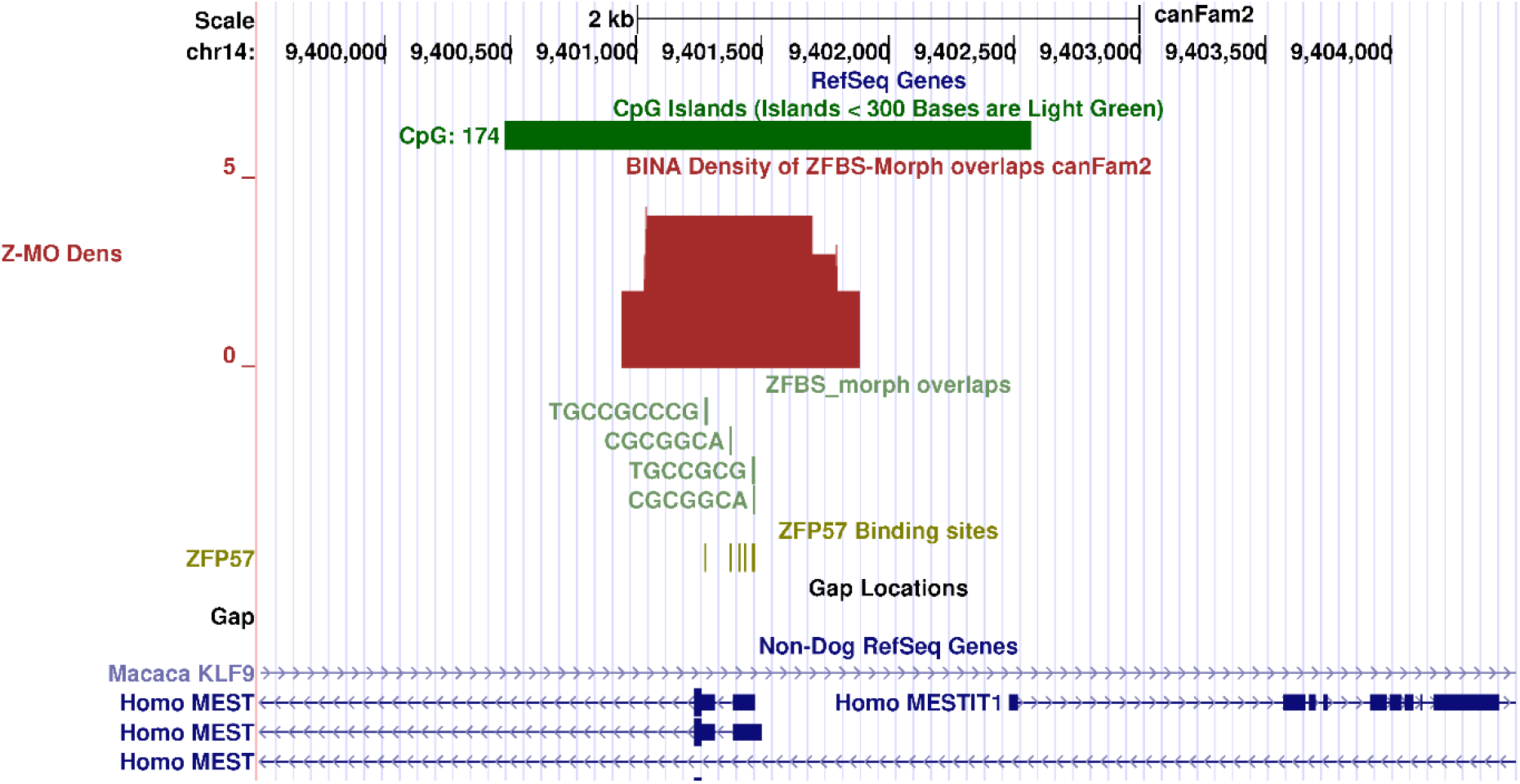
Candidate ICR for allele-specific expression of *MEST* isoform (*PEG1*) in Boxer genome. From top to bottom: the positions of CpG islands, in pack format; density-plot, in full format; ZFBS-morph overlaps, in pack format; ZFP57 binding sites, in dense format; and Non-Dog RefSeq Genes, in pack format. In the density-plot, the candidate ICR is marked by a robust peak in CpG174.

Functionally, in asexual reproduction (parthenogenesis), the absence of paternally imprinted genes often causes embryonic lethality [20]. Nonetheless, detailed gene targeting experiments facilitated introducing a mutation in *Peg1* to create viable and fertile female mice [25]. Due to the loss of *Peg1*, these females displayed unusual behavior [25]. Their response is relevant to studies of canines. Initially, the mice appeared normal: they delivered at term with a normal pregnancy rate [25]. They also showed an initial normal investigative behavior towards their pups. However, they did not demonstrate the expected maternal response to feeding and nurturing [25]. Furthermore, viable females displayed impaired placentophagia: a partly nutritional behavior observed in many mammals [25].

### A candidate ICR in deduced *PLAGL1* locus in Boxer

According to nucleotide databases, PLAGL1 related proteins are translated from several mRNA isoforms. Members of the PLAGL1 family bind DNA to regulate gene expression [26]. Human *PLAGL1* locus encompasses an intronic CpG island that includes the initiation site for a maternally expressed transcript known as *LOT1* or *ZAC1* [27]. Occasionally, *ZAC1* is referred to as *PLAGL1* [27, 28]. In addition to *ZAC1*, human DNA includes another imprinted intronic gene (*HYMAI*) transcribed to produce a noncoding RNA [29]. With respect to Non-Dog RefSeq genes, Boxer DNA includes several *PLAGL1* transcripts (Fig. 2). As reported for human and mouse [27, 28], Boxer locus also contains an intronic CpG island (CpG137). This island includes the TSS of both *HYMAI* and *ZAC1*, referred to as PLAGL*1* (Fig. 2). Density plots include a robust peak in CpG137. Therefore, the strategy correctly identified a candidate ICR for imprinted expression of both *ZAC1* and *HYMAI* in Boxer DNA (Fig. 2).

**Figure 2.**
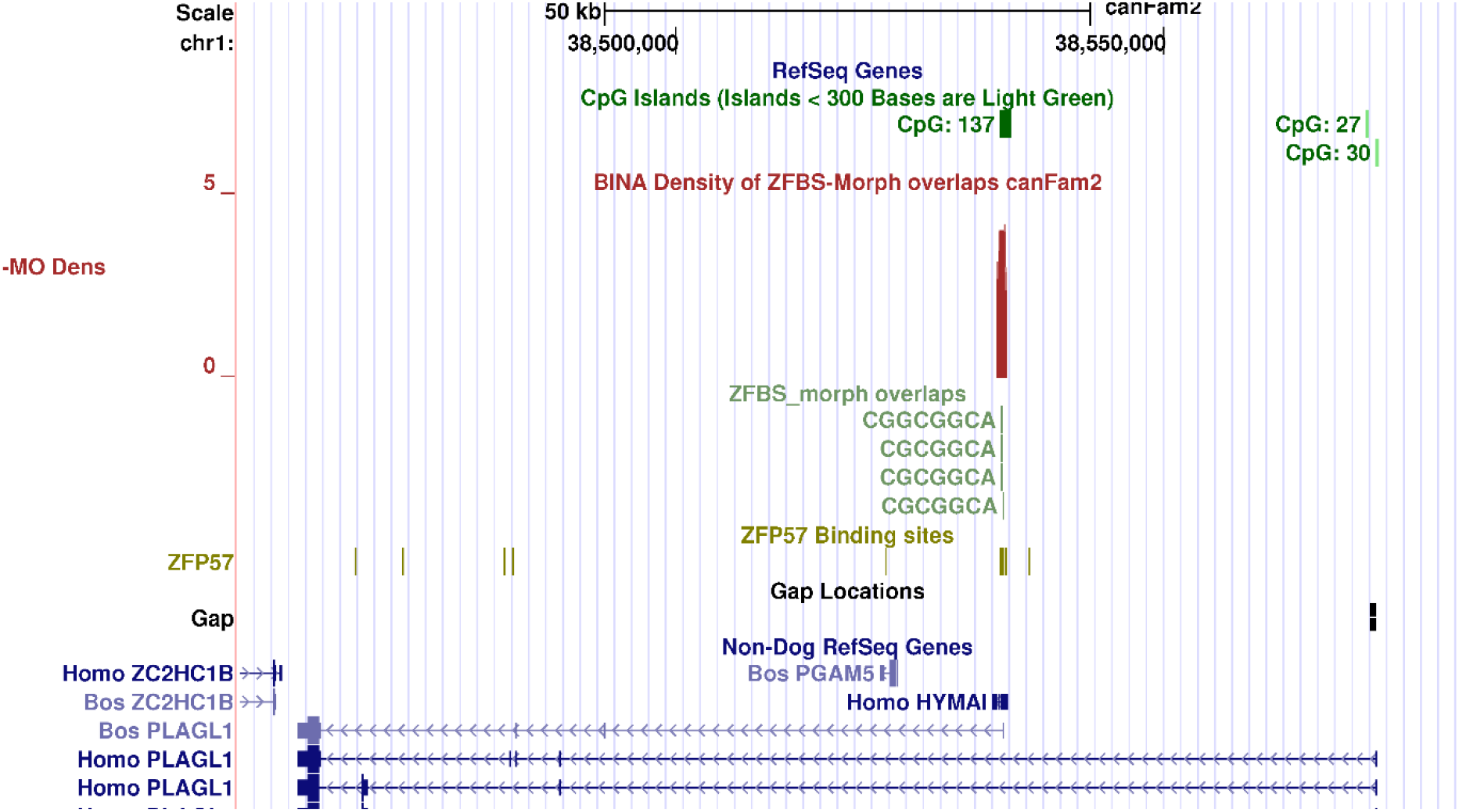
Candidate ICR for allele-specific expression of *PLAGL1* isoform (*ZAC1*) in Boxer genome. From top to bottom: the positions of CpG islands; density plot; ZFBS-morph overlap; ZFP57 binding sites; and Non-Dog RefSeq Genes. The candidate ICR is marked by a robust peak in CpG137.

### A candidate ICR in deduced *INPP5F* locus in Boxer

Also known as SAC2, INPP5F is a member of the inositol polyphosphate-5-phosphatase family of enzymes [30]. In both mouse and human, nearly all INPP5F transcripts are produced biallelically from a relatively long gene consisting of many exons and introns [31]. In combination with chromosome anomalies in mouse, tissue-specific microarray screening identified an imprinted transcript (*Inpp5f_v2*) expressed from the paternal allele in the brain [31]. This transcript has a unique alternative first exon [31]. Its TSS is within an intronic CpG island that includes the ICR for repressing *Inpp5f_v2* expression from the maternal allele [31]. Similarly, with respect to Non-Dog RefSeq Genes, Boxer genome includes an intronic CpG island (CpG153). In density plots, a peak is evident within that island. In the context of *Inpp5f_v2* TSS in mouse [31], the peak in CpG153 correctly marks the position of a candidate ICR for imprinted *INPP5F_V2* expression in Boxer DNA (Fig. 3).

**Figure 3.**
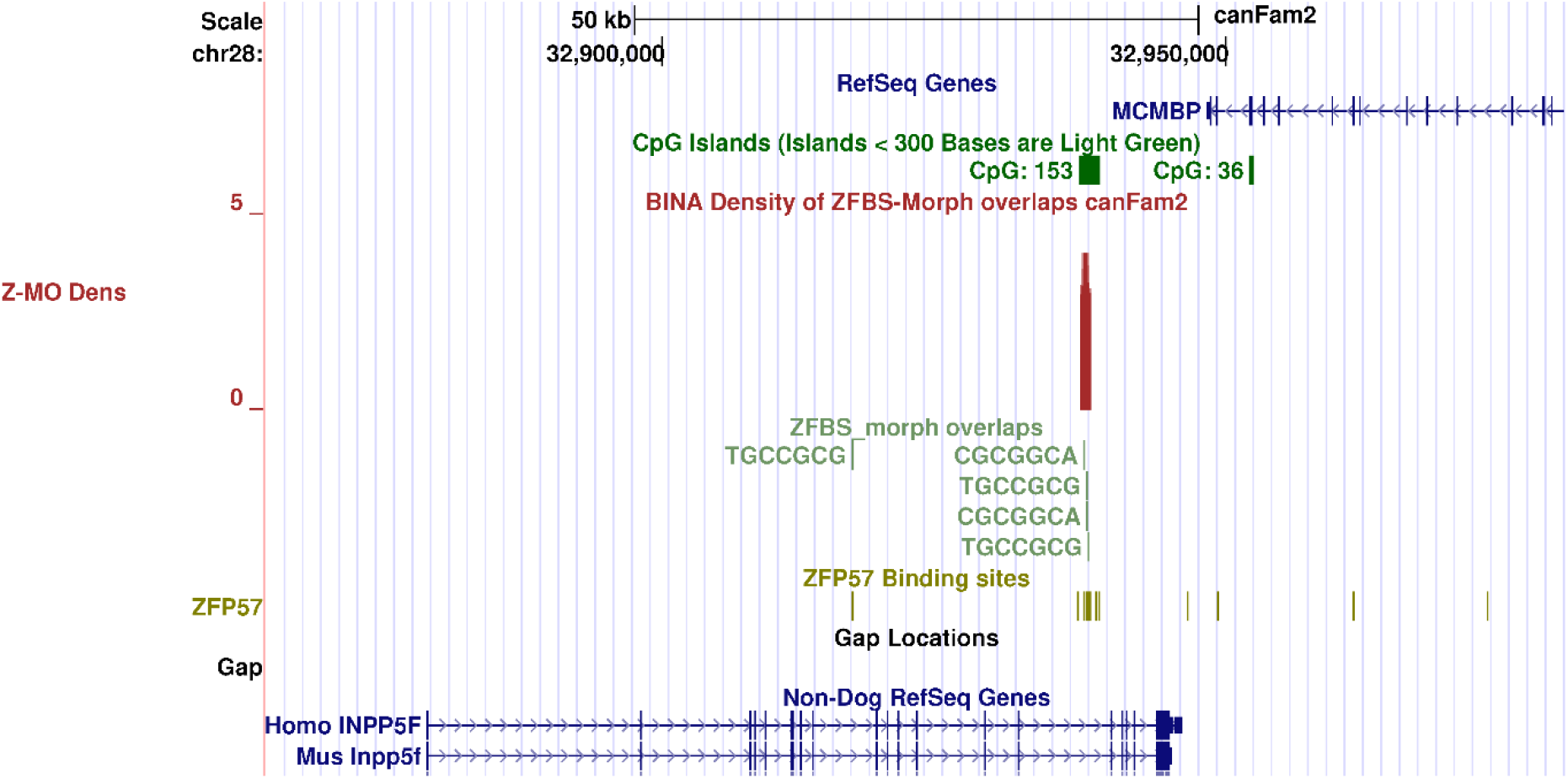
Candidate ICR for allele-specific expression of *INPP5F_V2* in Boxer genome. From top to bottom: the positions of CpG islands; density plot; ZFBS-morph overlaps; ZFP57 binding sites; and Non-Dog RefSeq Genes. The candidate ICR is marked by a robust peak in CpG153. In the context of studies of mouse and human, the ICR is intronic and at a correct position.

### A candidate ICR in deduced *ZIM2*/*PEG3* locus in Boxer

In mouse, *Peg3* regulates fetal growth and maternal behavior towards her pups [32]. The *Peg3* imprinted domain includes several genes expressed from the paternal allele and a few expressed from the maternal allele [33]. In mouse, a single CpG-rich gDMR/ICR regulates parent-of-origin-specific expression of the entire *Peg3* imprinted domain [33]. In mouse, the TSSs of *Zim*2 and *Peg3* are at differing positions; in human, transcription start sites of *ZIM*2 and *PEG3* coincide [33]. As in mouse, human *PEG3* is imprinted on the maternal allele [33].

As observed in mouse [33], in Boxer genome, a relatively long DNA segment encompasses several CpG islands. With respect to human RefSeq Genes, Boxer includes a gene annotated as Homo ZIM2 (Fig. 3). The bioinformatics strategy identified a candidate ICR for allele-specific expression of *PEG3/ZIM*2 in Boxer. In density-plots the candidate ICR is defined by a peak in the CpG island that encompasses the first exon of the transcript annotated as ZIM2 (Fig. 3). Therefore, this peak correctly located a candidate ICR for the imprinted expression of *ZIM2*/*PEG3* in Boxer DNA.

### A candidate ICR in deduced complex *GNAS* locus in Boxer

*GNAS* locus encompasses several related transcripts expressed from different promoters [34]. Its complexity stems from pre-mRNA splicing producing related transcripts [34]. Among *GNAS* transcripts, *Gαs* is biallelically produced, *XLαs* is expressed from the paternal allele [34]. Both *Gαs* and *XLas* specify related G-proteins with shared and distinguishable properties [35]. G-protein-coupled receptors (GPCRs) respond to extracellular inputs to relay information across outer cellular membrane to evoke proper physiological outcomes [36]. Downstream effects of a GPCR depends on which G protein type(s) it is coupled with [37].

A principal ICR regulates allele-specific expression of several *GNAS* transcripts [38]. Since *XLas* encodes a G-protein, it is essential to identify the ICR that regulates its parent-of-origin-specific expression. In Boxer DNA, a search located the complex *GNAS* locus with respect to human RefSeq Genes (Fig. 5). Examination of density plots revealed a peak in a CpG island. Since this peak encompasses *XLas* TSS, it offers a candidate principle ICR for the complex *GNAS* locus in Boxer DNA (Fig. 5).

**Figure 4.**
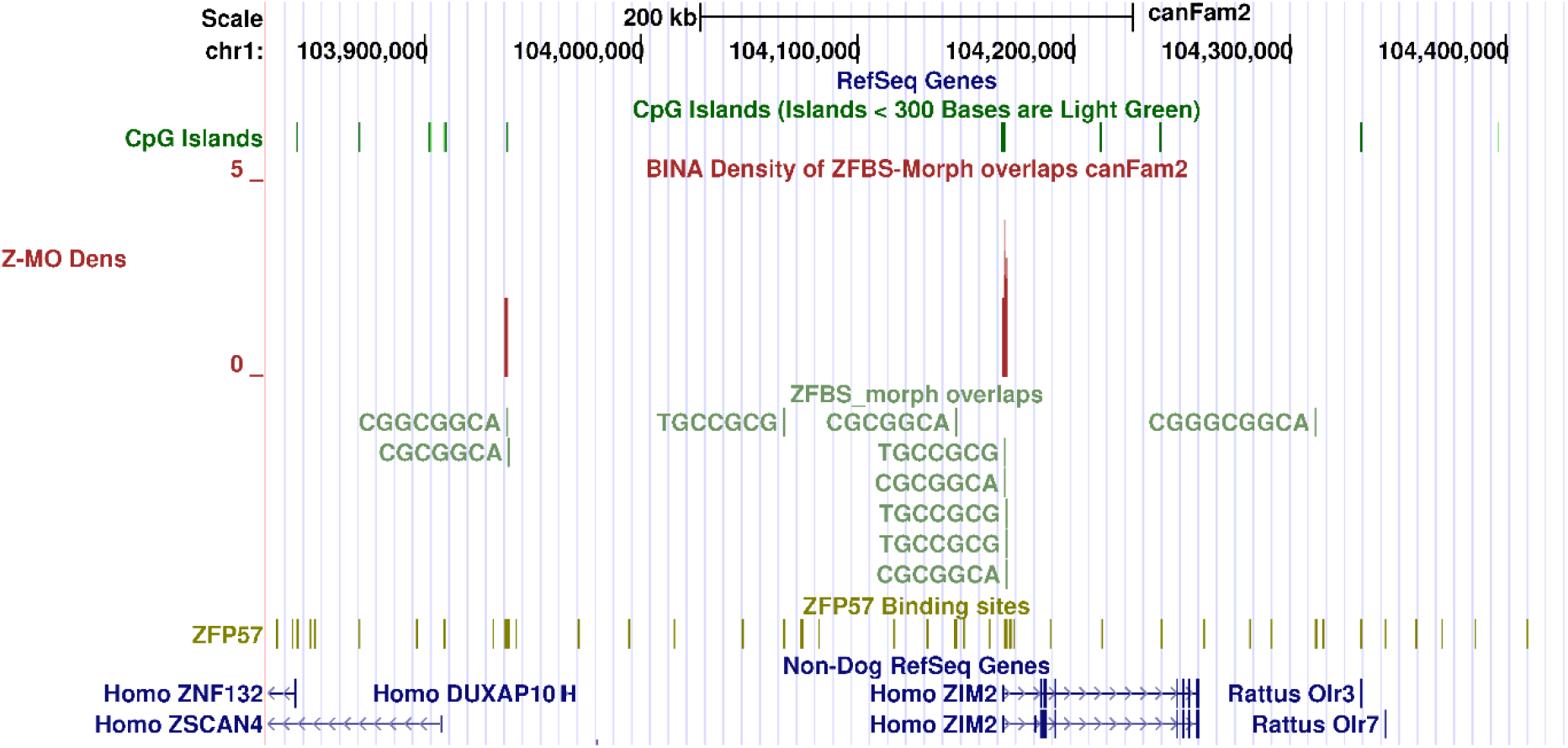
Candidate ICR for allele specific expression of *PEG3*/*ZIM2* in Boxer genome. From top to bottom: the positions of CpG islands; density plot; ZFBS-morph overlaps; ZFP57 binding sites; and Non-Dog RefSeq Genes. The robust peak marks the candidate ICR location. In the context of studies of human, candidate ICR position is correct.

**Figure 5.**
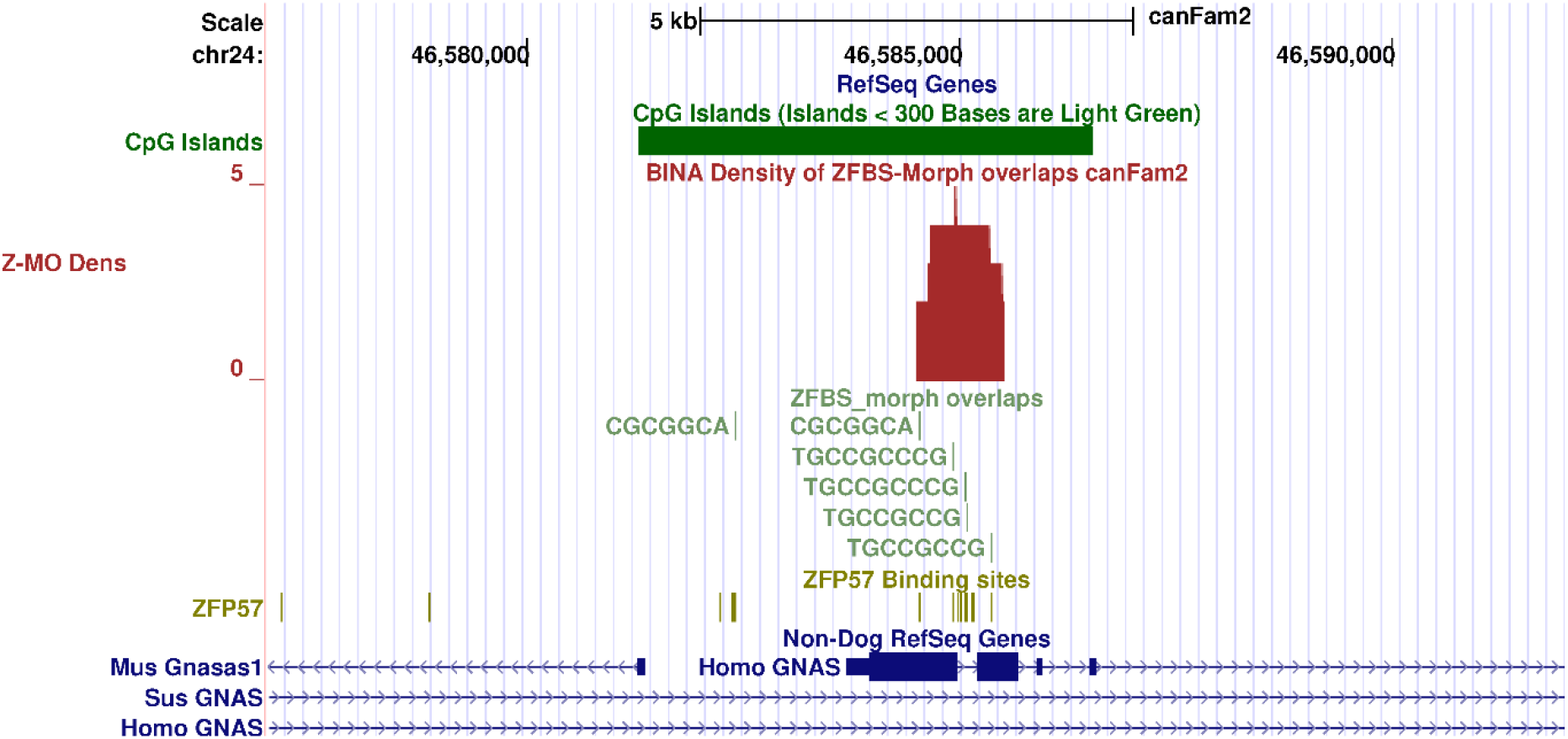
Candidate for principal ICR in the *GNAS* complex locus in Boxer genome. From top to bottom: the positions of CpG islands; density plot; ZFBS-morph overlaps; ZFP57 binding sites; and Non-Dog RefSeq Genes. The robust peak in marks the likely position of a candidate for principal ICR in *GNAS* complex locus in Boxer.

### A candidate ICR in *IGF2R* locus in Boxer

In mouse, the *Igf2r* imprinted domain encompasses at least two DMRs [39]. Even though the DMR at the *Igf2r* promoter is methylated on the repressed paternal allele, it does not exhibit expected characteristics of an ICR [39]. The other DMR is a domain-wide ICR [39]. In mouse, the ICR is in the second *Igf2r* intron and encompassed by a CpG island [40]. The intronic DNA also includes the TSS of an imprinted noncoding RNA gene known as *Airn* [39]. With respect to *Igf2r, Airn* is in an antisense orientation [39]. Evidence suggested a role for *Airn* in regulating imprinted *Igf2r* expression [39]. However, in dog, *IGF2R* was expressed monoallelically even in the absence of *AIRN* [41]. The bioinformatics strategy predicted a candidate ICR in the second intron of *IGF2R* in Boxer DNA (Fig. 4). This ICR is defined by a peak in density plots. Its position agrees with the intronic ICR position in mouse [40].

### Genomic imprinting of *H19* – *IGF2* domain in *Canis familiaris*

In canine, sequences upstream of *H19* encompass reiterated DNA [42]. They include two direct repeats with partial internal repeats. The domain encompasses a putative differentially methylated ICR with likely paternal transmission [42]. Previously, the bioinformatics strategy correctly identified the ICR of the *IGF2*–*H19* imprinted domain in mouse [17], human [43], and *Bos taurus* [44]. However, since the strategy relies on detecting clusters of ZFBS-morph overlaps, it did not predict a candidate ICR in Boxer DNA (Fig. 7). Upstream of *H19*, Boxer genome encompasses three ZFBS-morph overlaps, and several hexameric sequences that after methylation, bind ZFP57 (Fig. 7). In mouse ESCs, binding of ZFP57 to methylated hexameric sites in ICRs was required to maintain allele-specific repression of imprinted genes [8]. Therefore, an intriguing question is whether ZFP57 also contributes to the regulation of the *IGF2*–*H19* domain in canine DNA.

**Figure 6.**
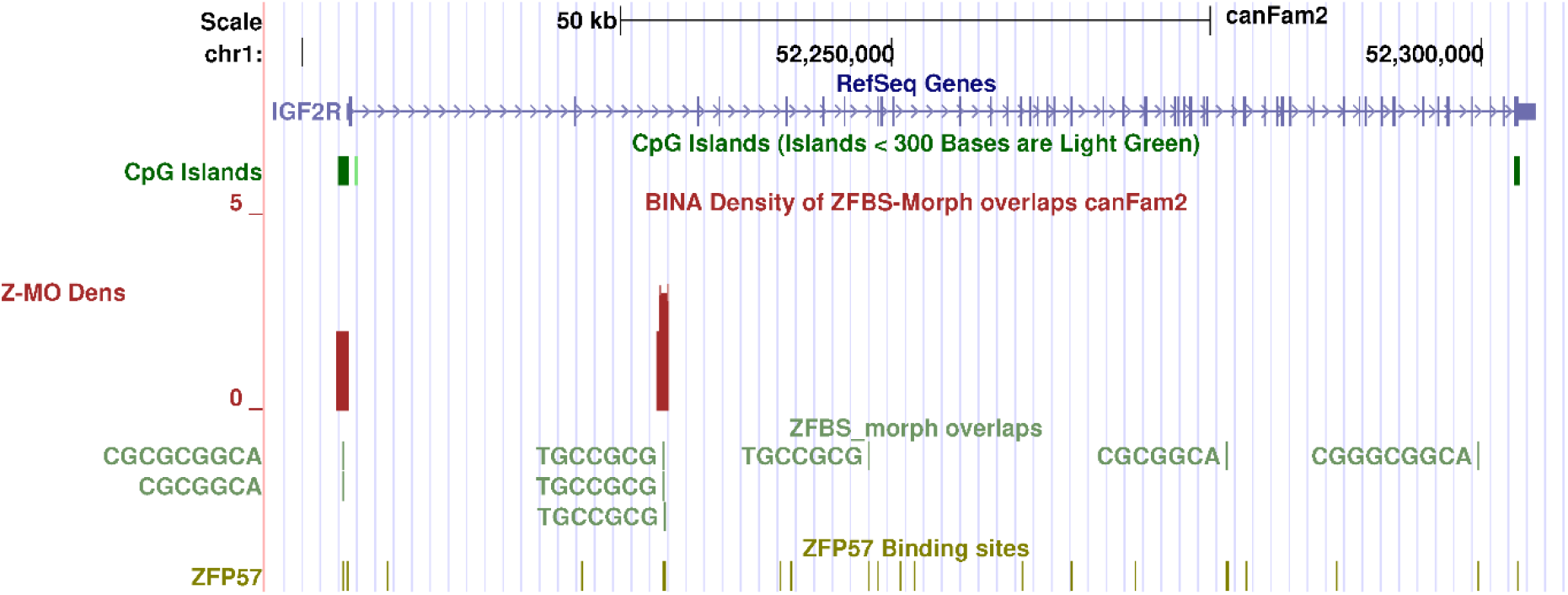
Candidate ICR for allele specific expression of *IGF2R* in Boxer genome. From top to bottom: the positions of CpG islands; density plot; ZFBS-morph overlaps; ZFP57 binding sites; and Non-Dog RefSeq Genes. The candidate ICR is intronic and marked by a robust peak. In the context of studies of mouse, the ICR is at correct intronic position in Boxer DNA. However, in contrast to mouse, in Boxer, the predicted ICR is not within a CpG island.

**Figure 7.**
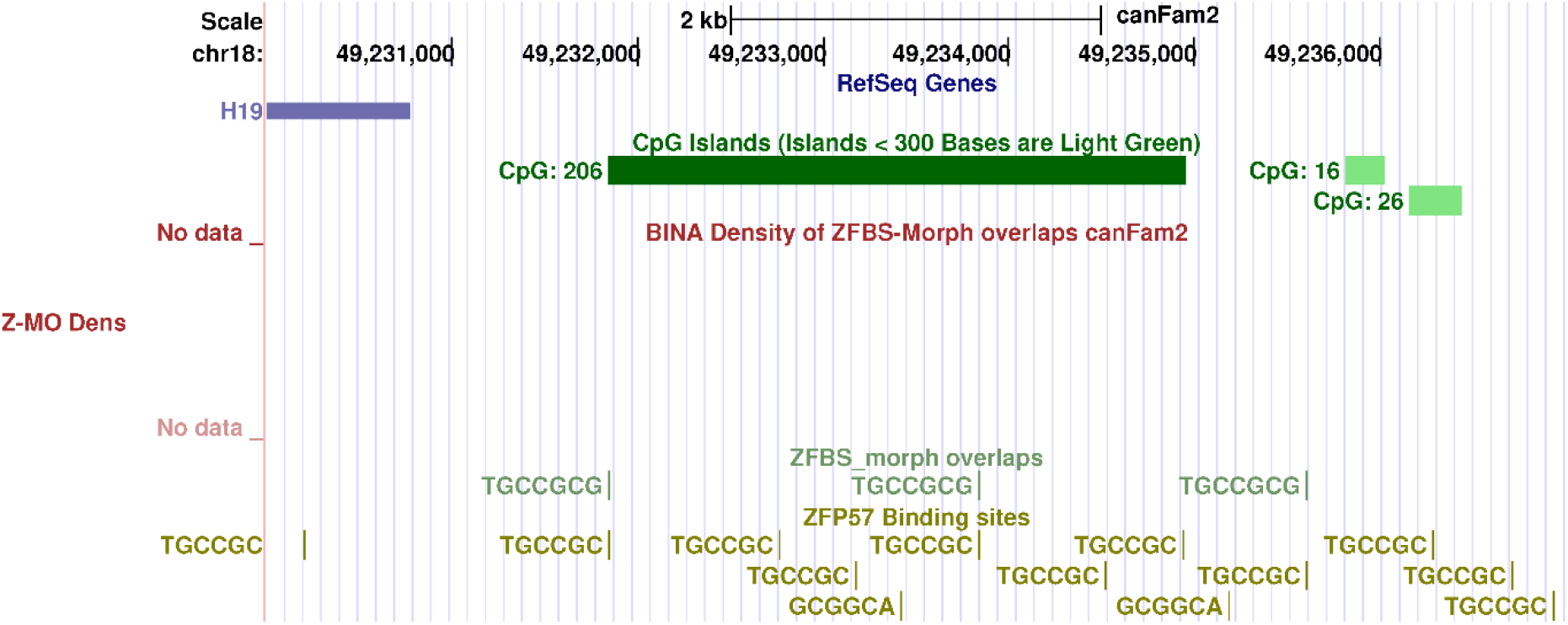
*H19* locus in Boxer genome. From top to bottom: the positions of *H19*, CpG islands, ZFBS-morph overlaps, and hexameric sequence. Upstream *H19*, the strategy did not find a candidate ICR for regulating gene expression in Boxer DNA. In the region, sequence analyses located 3 ZFBS-morph overlaps, and several hexamers that after methylation, would bind ZFP57.

## DISCUSSION

As in other mammals, imprinted genes are likely to play critical roles in embryogenesis and fetal development in *Canis familiaris* [45]. However, only a few studies have addressed genomic imprinting in dog [42, 45, 46]. Therefore, locating candidate ICRs in Boxer genome could help with identifying genes with likely impact on canids embryonic development, behavior, and physiology. Previously, our bioinformatics strategy located a substantial fraction of known ICRs in mouse, human, and Bos taurus [17, 43, 44]. Since the strategy is genome-wide, it facilitates discovering candidate ICRs for novel potential imprinted genes [17, 43, 44, 47].

In Boxer, with respect to Non-Dog RefSeq Genes, the strategy predicted candidate ICRs for allele-specific expression of *PEG1*/*MEST* isoform, *PLAGL1*/*ZAC1* isoform, *INPP5F_V2* isoform, *ZIM2*/*PEG3, IGF2R*, and *GNAS XLαs* (Figs, 1-6). Why these loci could be of significance to studies of *Canis familiaris*? Because, they include genes whose products are likely to impact embryogenesis and development in dogs, as observed in mouse [45]. For example: in mouse, the *Mest* isoform *Peg1* influences maternal response to her newly born pups [25]. During mouse embryogenesis, *Zac1* is transcribed in the progenitor/stem cells of several tissues including neuronal and skeletal muscle [48]. The encoded protein (ZAC1) plays key roles in the development of the pituitary [49]. Furthermore, ZAC1 regulates cell-specific expression of *PAC1R* [50]. The encoded protein (PAC1R) is a G protein-coupled receptor. In response to PACAP, PAC1R transmits signals to the neuroendocrine, endocrine, and nervous systems [50, 51]. In dog ileal circular muscle, PACAP operates as a neurotransmitter [52]. In anesthetized dogs, PAC1R functions in adrenal catecholamine secretion induced by vasoactive intestinal polypeptide (VIP) and PACAP [53].

In mouse, *Inpp5f_v2* is expressed in the brain [31]. Mutations in paternal *Peg3* caused maternal behavioral defects in nurturing and low offspring survival [32]. in carrier females, loss of paternal *Peg3* impaired milk letdown [32]. In part, poor behavioral manifestations could be due to anomalies in oxytocin signally pathway in *Peg3* deficient females [32]. Known as “the love hormone” [54], oxytocin mediates dogs’ human-directed social behavior [55].

In the *Gnas* complex locus, one of the imprinted transcripts (*XLαs*) encodes a G-protein that functions in signaling pathways by several GPCRs [56]. *XLαs* expression was detected in distinct regions of the midbrain, the hindbrain, the noradrenergic system of the brain, and in neuroendocrine tissues [34, 57]. Furthermore, XL*αs* is crucial for early postnatal adaptation to feeding and survival [34]. Lastly, IGF2R (insulin-like growth factor 2 receptor) facilitates IGF2 endocytosis [45]. In murine, developmental anomalies result from *Igf2r* mutations in the maternal allele [58]. Thus, Overall, the bioinformatics strategy has identified candidate ICRs for several genes with likely impact on canine development, behavior, and physiology. Furthermore, since the strategy was applied genome-wide, researchers could discover candidate ICRs for novel imprinted genes in canines.

## METHODS

### Marking the genomic positions of ZFP57 binding site and the ZFBS-morph overlaps

The UCSC genome browser offers a link for downloading the nucleotide sequence of the build CanFam2 of Boxer genome. A Perl script identified the genomic positions of the hexameric sequence TGCCGC that after CpG methylation binds ZFP57 [8]. Another Perl script located genomic positions of ZFBS-morph overlaps. UNIX subroutines combined both outputs to create a file suitable for upload on the UCSC genome browser to create two custom tracks.

### Creating plots of ZFBS-morph overlaps density

A Perl script opened the file containing the genomic positions of ZFBS-morph overlaps in a specified chromosome. The script scanned the file to count and to report the number of ZFBS-morph overlaps in a sliding widow consisting of 850-bases. To remove background noise, the script ignored isolated occurrences. The outputs consisted of peaks covering 2 or more ZFBS-morph overlaps. Peak positions were determined with respect to the midpoint of each window. The length of the window was selected by trial and error [14]. Large windows produced false peaks; small windows yielded peaks with a spiky appearance. Subsequently, a UNIX subroutine combined and tailored the outputs for display as a custom track on the UCSC genome browser. Shown figures were downloaded as PDF produced on the genome browser; the goal was to display the custom tracks with respect to underlying genomic positions in Boxer DNA. While creating the datasets, CanFam2 was the latest available build of Boxer genome.

## Author contributions

P. Wyss wrote the Perl scripts. M. Bina analyzed Boxer sequences and wrote the manuscript.

## Acknowledgment

We thank Maya Afilalo for checking the manuscript.

## Data availability

Datasets can be accessed for download from Purdue University Research Repository (PURR).

- The positions of ZFBS and ZFBS-morph overlaps in the build canFam2 of the dog genome https://purr.purdue.edu/publications/3519/1
- Density of ZFBS-morph overlaps in the build canFam2 of the dog genome https://purr.purdue.edu/publications/3520/1

## Ethics declarations

The research did not include human or animal subjects.

